# Crossmodal attention develops in the first year of life: Cortical signatures of tactile to visual exogenous spatial cuing at 8 but not 5 months of age

**DOI:** 10.64898/2025.12.03.691887

**Authors:** Giulia Orioli, Rhiannon L. Thomas, Jannath Begum Ali, Luke Mason, Jose L. van Velzen, Andrew J. Bremner

## Abstract

Adults are able to orient their attention between different (often fleeting) sensory cues that present to different senses (“crossmodal attention”), but we cannot assume that this is the case in early life. Here we report the findings of a study in which we probed the presence of tactile to visual crossmodal links in spatial attention using event related potentials (ERP) gathered from the scalp electroencephalograms (EEG) of 5-(n = 19) and 8-month-old (n = 19) infants. Whilst recording EEG, we presented 5- and 8month-old participants with vibrotactile stimuli on one of their hands, followed by visual stimuli on the same (Congruent) or the opposite (Incongruent) hand, or no touch at all (No probe). During the presentation of these stimuli the infants’ eyes were oriented centrally. We subtracted No probe trials from Congruent and Incongruent trials to yield visual evoked potentials (VEPs) free from the influence of prior somatosensory processing. Comparably to prior findings in adults, 8-month-olds’ first negative component of the VEP was enhanced in the hemisphere ipsilateral to the probe when the visual probe was spatially congruent with the tactile cue. No crossmodal effect was observed in the 5-month-olds, indicating developments in the crossmodal coordination of attention in the first year of life.

## INTRODUCTION

Adults are able to orient their attention between different (often fleeting) sensory cues that present to different senses (“crossmodal attention”; Spence & Driver, 2004), but we cannot assume that this is the case in early life. Despite considerable research into the early development of multisensory perception (see Bremner et al., 2012), and selective attention to multisensory stimuli (e.g. Bahrick et al., 2004, 2020), we know little about the ontogeny of selective attentional shifts *between different sense modalities*. If such abilities undergo significant early development, there are wide-ranging implications for how infants experience their multisensory environments. Here we report the findings of a study in which we probed tactile to visual crossmodal links in spatial attention using event related potentials (ERP) gathered from infants’ scalp electroencephalograms (EEG). Eight-, but not 5-month-old infants showed an influence of a tactile cue on subsequent visual processing at the same location.

Soon after birth infants show impressive competencies in selecting multisensory information (e.g. Bahrick et al., 2020; Bahrick & Lickliter, 2000; Bremner et al., 2012). Newborns quickly learn associative links between the senses (Orioli et al., 2018; Schaal & Durand, 2012; Slater et al., 1999) and young infants capitalise on information in one sense to enhance processing in another (Bahrick & Lickliter, 2000; Durand et al., 2020; Leleu et al., 2020; Orioli et al., 2023), raising the question of how such associations and crossmodal interactions are achieved. One possibility is that young infants are able to engage crossmodal attention in prioritising the processing of information occurring at the same (or nearby) time supramodally.

Overt crossmodal orienting responses where infants orient one sensory system (e.g. the eyes) towards a stimulus perceived through another sense (e.g., towards a sound or a touch) provide important clues to the origins of crossmodal attention. In the case of visual-auditory orienting, newborns are known to make saccades (Butterworth & Castillo, 1976; Wertheimer, 1961) and head movements (Clifton et al., 1981; Muir & Field, 1979) coordinated with a sound location. However, these early behaviours disappear in 3-month-olds to be replaced by more accurate and flexible orienting responses from 4-5 months (Muir et al., 1989). Evidence for neonatal crossmodal orienting to *touch* is scarcer. Moreau, Helfgott, Weinstein, and Milner (1978) have shown habituation of neonatal head turning to tactile stimuli. Later, Bremner, Mareschal, Lloyd-Fox, & Spence (2008) found evidence for accurate crossmodal orienting of the eyes to touches at 10 but not 6 months of age. Regardless, overt crossmodal orienting responses are not a transparent marker of the existence of crossmodal links in attention or vice versa. Adults show covert crossmodal attentional enhancement, independently of motor responses (Hunt & Kingstone, 2003b, 2003a; Spence & Driver, 1996).

A number of studies have used adaptations of the Posner cuing task (Posner, 1980) to investigate covert shifts of attention in infancy within a single sense modality. These studies have made use of infants’ tendency to orient their eyes towards targets, measuring the effect of valid and invalid cues presented prior to saccades on subsequent target-directed eye-movement latencies. As infants cannot be asked to maintain central fixation, overt cue orienting is avoided either by brief presentation (Johnson & Tucker, 1996) or by maintaining a salient fixation element while the cue is presented (Richards, 1987, 1997). Both facilitation and inhibition of return in covert visual cuing have been observed in infants by 6 months of age (Hood, 1993; Hood & Atkinson, 1993), with some indications that covert attention may emerge between 2 and 6 months of life (Johnson, 1994; Johnson et al., 1994; Johnson & Tucker, 1996).

Event-related potentials (ERPs, Richards, 2000) and steady-state visually evoked potential (SSVEP, Christodoulou et al., 2018) have also been used as indices of covert orienting. In a study of 4- to 6-month-old infants, Richards and colleagues (2000) demonstrated a larger contralateral, occipital P1 component in response to the valid trials (in contrast to the invalid and the control trials), suggesting that covert attention shifts may affect the early stages of visual processing. They also found larger presaccadic responses to the target when this appeared in the cued vs uncued location. More recently, Christodoulou, Leland, and Moore (2018) used steady-state visually evoked potentials (SSVEPs) to investigate covert shifts of visual attention in 4-month-old infants. Together these findings provide conspicuous evidence of the mechanisms of covert attention in young infants, but are limited to conditions where both cue and target are presented in the visual modality. Given that most of the stimulation provided to developing infants by the environment is conveyed across different senses, even prenatally, the focus of this study is on crossmodal covert attentional shifts.

Here, we set out to examine whether tactile stimuli presented to a single hand would cue enhancements of visual processing at the same location in human infants. Our study had two objectives. Firstly, we wanted to examine whether it is possible to measure crossmodal spatial orienting effects (between touch and vision) in preverbal infants, starting from 8 months of age. We chose this age group because: i) unimodal (visual) covert spatial orienting effects are established by this age (from 6 months, Richards, 2000) and, ii) the influence of a visual frame of reference on tactile orienting behaviour is established by this age (in 6-month-olds, Azañón et al., 2018; Begum Ali et al., 2015). After establishing that the 8-month-olds demonstrated an enhancement of visual processing at a location that had just been cued by a tactile cue, we next examined whether we could observe the development of this crossmodal cuing effect in the first postnatal year via a comparison to a new sample of 5-month-old infants. The 5month-old age group given the development of visual-tactile links in spatial behaviour up to 6 months of age (Azañón et al., 2018; Begum Ali et al., 2015)).

We based our task on a procedure carried out with adult participants (Kennett et al., 2001) and examined ERP signatures of links between vision and touch in exogenous spatial attention. When presenting their adult participants with non-predictive tactile cues to either hand, followed by visual probes shown in either the same or contralateral hemifields, Kennett et al. (2001) found a larger amplitude in the ipsilateral occipital N1 when the tactile cue and the visual probe were congruent, i.e., presented on the same side of the body, as opposed to opposite sides, demonstrating a crossmodal cuing effect of tactile cues on visual evoked potentials. Here we presented three conditions: i) Congruent cue-probe condition in which a short vibrotactile cue was presented on either the left or the right hand, followed by a visual probe on the same hand, ii) Incongruent cue-probe condition in which tactile cue and visual probe were presented on different hands, and iii) No probe condition in which the tactile cue only was presented. The No Probe condition was included to assess the presence of genuine crossmodal influences on visual processing rather than the persistence of tactile processing signatures bleeding over to visual scalp areas (the No probe condition was subtracted from each of the other conditions).

## METHOD

### Participants

The study involved two groups of 19 infants, predominantly Caucasian, aged 5 and 8 months. We initially recruited a group of 8-month-old infants (February 2014 - June 2016). Based on the results, suggesting differences in the Ipsilateral N1 in response to congruent and incongruent Cue-Probe pairings, we decided to recruit a group of younger infants, aged 5 months (June-July 2017 and April-December 2018), to investigate when such difference emerges during development. The study was not preregistered.

Previous studies measuring somatosensory evoked potentials in infants required 15 participants to uncovered significant differences between conditions (Rigato et al., 2019). Here, we increased the sample size to 19 to improve statistical power: a two-tailed t-tests with *α* = 0.05 and power = 0.85 requires 19 participants if a medium effect size (*d* = 0.7) is expected (sample size calculated using G-Power, Faul et al., 2007)).

The 5-month-old infants (12 female) were aged on average 21.77 weeks (SD = 1.68 weeks, range: 19.14-24 weeks). A further 8 infants participated, but were excluded due to fussiness or excessive movement (2), experimental error (1), or because they did not contribute enough artefact-free trials (7 per trial type, 5). The 8-month-old infants (10 female, 8 male, sex for one participant not recorded) were aged on average 32.4 weeks (SD = 1.26 weeks, range: 30.14-36 weeks). The number of infants who participated but were excluded infants and the reasons for exclusion cannot be reported due to data loss (caused by a hard drive failure).

Participants were recruited from the [blinded] database and testing took place at the [blinded] when the infants were awake and alert. Parents were informed about the procedure and provided informed consent for their child’s participation. Ethical approval was gained from the Institutional Ethics Committee [blinded], and the study was conducted in accordance with the Declaration of Helsinki. Participants received a small gift in return for their contribution.

### Design

The study involved the presentation of tactile cues and visual probes across the infants’ two hands. Different combinations of cues and probes contributed to the formation of 3 Cue-Probe conditions: Congruent Cue-Probe, Incongruent Cue-Probe and No Probe. Each condition included a tactile cue on one hand, followed either by a visual probe on the same hand (Congruent condition), or a visual probe on the opposite hand (Incongruent condition), or by no visual probe (No Probe condition). No-Probe trials were included as a control to ensure that any effects of condition on EEG responses to the visual probe were not influenced by purely somatosensory activity propagating to visual electrodes (see EEG Recording and Analyses). We recorded the infants’ brain activity throughout the study using electroencephalography (EEG) and analysed the visual evoked potentials (VEPs) measured in response to the visual stimuli. For both the Congruent and Incongruent conditions, we calculated difference waveforms between the trials with and without the visual probe to measure and compare purely the visual responses and ensure that the tactile components, common across all trials, were removed by the subtraction.

The cue and probe stimuli were presented across the left and the right hands, leading to a total of 6 types of trials (cue and probe on the left hand - Congruent, cue and probe on the right hand - Congruent, cue on the left hand and probe on the right hand - Incongruent, cue on the right hand and probe on the left hand - Incongruent, cue on the left hand and no probe - No probe, cue on the right hand and no probe - No probe). Trials were presented in sets of 4. In each set, 4 trial types were randomly picked from the 6 trial types available and presented in random order. The trial presentation was continued until the infants’ attention lasted and up to 60 sets of 4 trials.

### Procedure, Stimuli and Apparatus

Each infant sat on their parent’s lap on a chair in front of a small table in a dimly lit room with black curtaining covering the walls. Tactors (custom-built voice coil tactile stimulators, PsyAl, https://www.psyal.co.uk/) were placed in each of the infant’s hands, secured using self-adherent bandage and covered with white cotton mittens. An LED was attached to each mitten and positioned on the back of the infants’ hands, so that the light could be visible. The experimenter sat in front of the infant and held their wrists as still as possible above the table, about 20 cm apart from each other.

The experimenter attracted the infant’s attention to her face, singing and playing games such as “peek-a-boo”. When the infant engaged with the experimenter, a second experimenter triggered the presentation of the first set of 4 trials and, monitoring the infant’s behaviour from outside the experimental room, continued to trigger subsequent sets of 4 trials until the infant’s interest lasted.

Each trial began with a 100 ms tactile cue (a short vibration) on one of the hands. This was followed by a 50 ms inter-stimulus interval, followed by a 200 ms visual probe (a) on the same hand (Congruent condition) or the opposite hand (Incongruent condition), or by no visual probe (No Probe condition). We selected a short stimulus onset asynchrony (SOA) of 150 ms between tactile cue and visual probe in line with the SOA used by Kennett et al. (2001) and in light of the facilitation effects in covert visual orienting that have been observed at these short SOAs in younger (3- and 4-month-old) infants (e.g. (Xie & Richards, 2017)). The visual probe (or its absence) was followed by an inter-trial interval randomly lasting between 800 and 1200 ms (Fig. 1). The tactors (driven by a 220 Hz sine wave) and the LEDs were controlled with EPRIME 1.1 (Psychology Software Tools, Pittsburgh, PA). The noise made by the tactors was covered by the experimenter’s voice throughout the experiment. The infants’ looking behaviour was recorded via a single infrared camera located in the left corner of the room.

**Figure 1.**
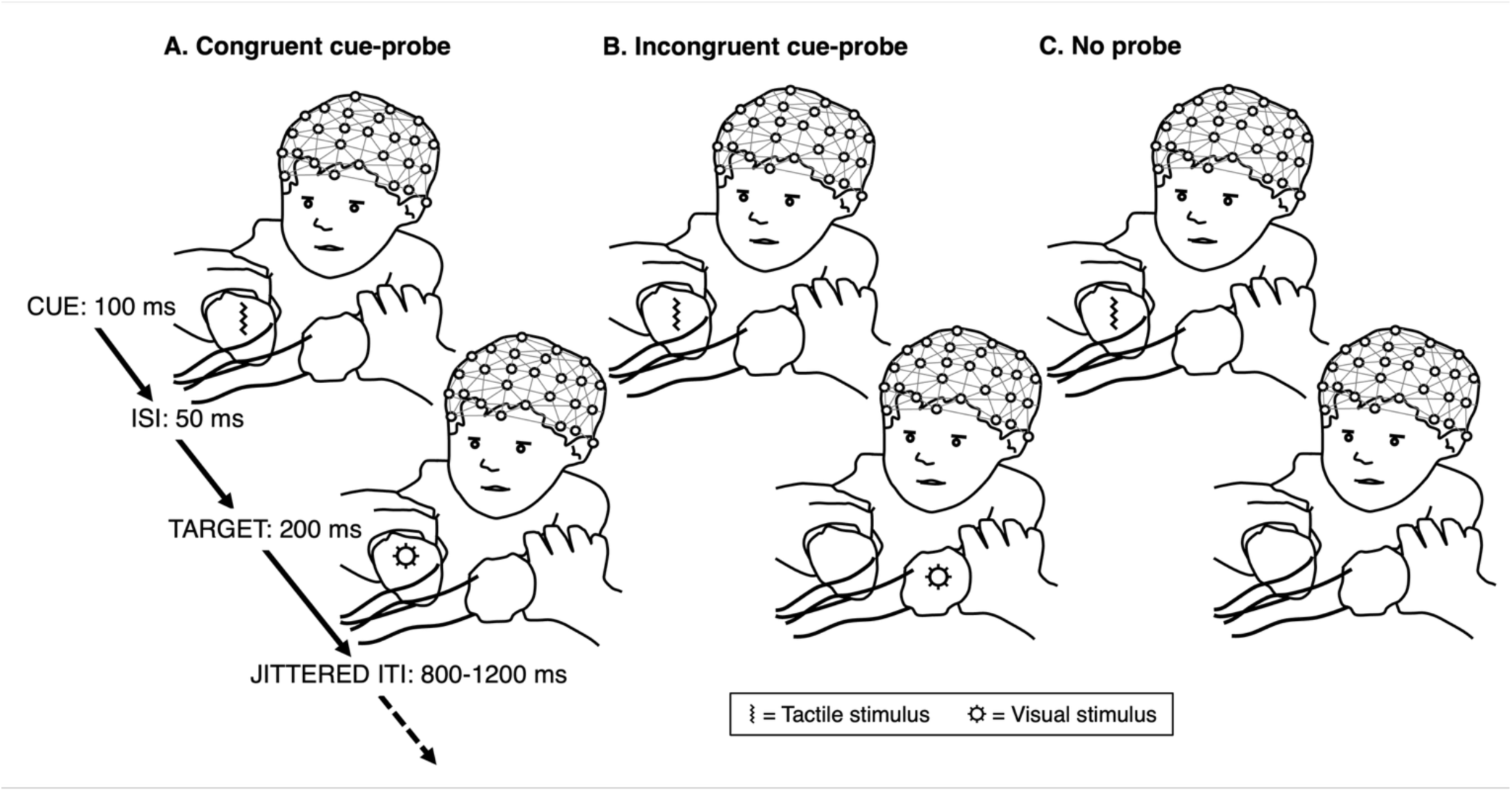
Experimental paradigm.

### EEG recording and analyses

Participants’ electrical brain activity was continuously recorded using a Hydrocel Geodesic Sensor Net (Electrical Geodesics Inc., Eugene, Oregon), consisting of 128 silver-silver chloride electrodes evenly distributed across the scalp and referenced to the vertex. The potential was amplified with a 0.1 to 100 Hz band-pass and digitized at 500 Hz sampling rate (Hoehl & Wahl, 2012). The raw data were processed offline using NetStation 4.5.1 (Electrical Geodesics Inc., Eugene, Oregon). Continuous EEG data were high-pass filtered at 0.3 Hz and low-pass filtered at 30 Hz using digital elliptical filtering (Hoehl & Wahl, 2012). They were segmented in epochs from 250 ms before the visual stimulus onset to 700 ms after it and baseline-corrected to the average amplitude of the 150 ms preceding the visual stimulus onset (i.e., from the onset of the tactile cue). Epochs containing movement artefacts or more than 12 bad electrodes (Hoehl & Wahl, 2012) were visually detected and rejected. Bad electrodes were interpolated trial-by-trial using spherical interpolation of neighbouring channel values. Video recordings of the session were coded offline to identify and exclude trials where the participant was distracted or was looking at their hands rather than the experimenter (for the 5-month-old group, 16/19 videos were available: removing the 3 participants whose videos where not available did not change the results, see Supplementary Fig. SI1 and Supplementary Table SI1). Artefact free trials (both age groups: M = 28 trials per trial type, see Supplementary Table SI2) were re-referenced to the average potential over the scalp, then individual averages were calculated. We next computed the difference waveform between the responses recorded in the Cue-Probe trials and those recorded in the No Probe trials.

To identify the electrode clusters for the analyses, we averaged the difference waves and inspected the topographic maps (using FieldTrip, Oostenveld et al., 2011; Popov et al., 2018) representing the scalp distribution of the electrical activity (Luck & Gaspelin, 2017), confirming the presence of hotspots in the regions surrounding O1 and O2 in the 10-20 system. Next, we visually inspected the recordings from the electrodes within these areas and isolated, for each hemisphere, the electrode cluster showing the most pronounced VEP components (Luck & Gaspelin, 2017): for the 5month-old group, 59, 60, 66 (left hemisphere) and 84, 85, 91 (right hemisphere); for the 7-month-old group, 66, 70, 71 (left hemisphere) and 76, 83, 84 (right hemisphere).

For the mean amplitude analyses, we selected the latencies of interest for the P1 and N1 components using a collapsed localisers approach (Luck & Gaspelin, 2017), which reduces the chance of biased measurements by defining component latency based on the average waveform across Cue-Probe conditions.

## RESULTS

We identified a positive VEP component (P1), followed by a negative VEP component (N1) both ipsilaterally and contralaterally to the visual stimulus. The latencies of these components were slightly different depending on whether they were recorded over ipsilateral or contralateral hemispheres. Because VEP components change significantly in amplitude and latency during the first year of life (De Haan, 2013; Webb et al., 2005), we treated the responses of the 8- and the 5-month-old infants separately.

### 8-month-olds

We compared the mean individual amplitude of the VEPs in the Congruent vs Incongruent Cue-Probe conditions in the P1 and N1 components. The latencies of these components were: Ipsilateral hemisphere, P1: 122-200 ms, peaking at 170 ms; N1: 202-258 ms, peaking at 240 ms; Contralateral hemisphere, P1: 114-198 ms, peaking at 166 ms; N1: 200-298 ms, peaking at 242 ms (Fig. 1a).

For each hemisphere and component, we ran a paired t-test to compare the mean individual amplitudes of the waveform between the two Cue-Probe conditions. The results show a significant difference in the mean individual amplitude of the VEP in the N1 component in the Ipsilateral hemisphere, larger for the Congruent than the Incongruent Cue-Probe condition, *t*(18) = −2.501, *p* = 0.022, *d_z_* = 0.574 (proportions of looking time were normally distributed, *D* = 0.165, *p* = 0.622). The mean individual amplitudes in the Congruent and Incongruent conditions were −2.734 μV (S.E. = 0.841 μV) and 0.346 μV (S.E. = 0.995 μV), respectively (Fig. 1b and Table 1).

**Figure 2.**
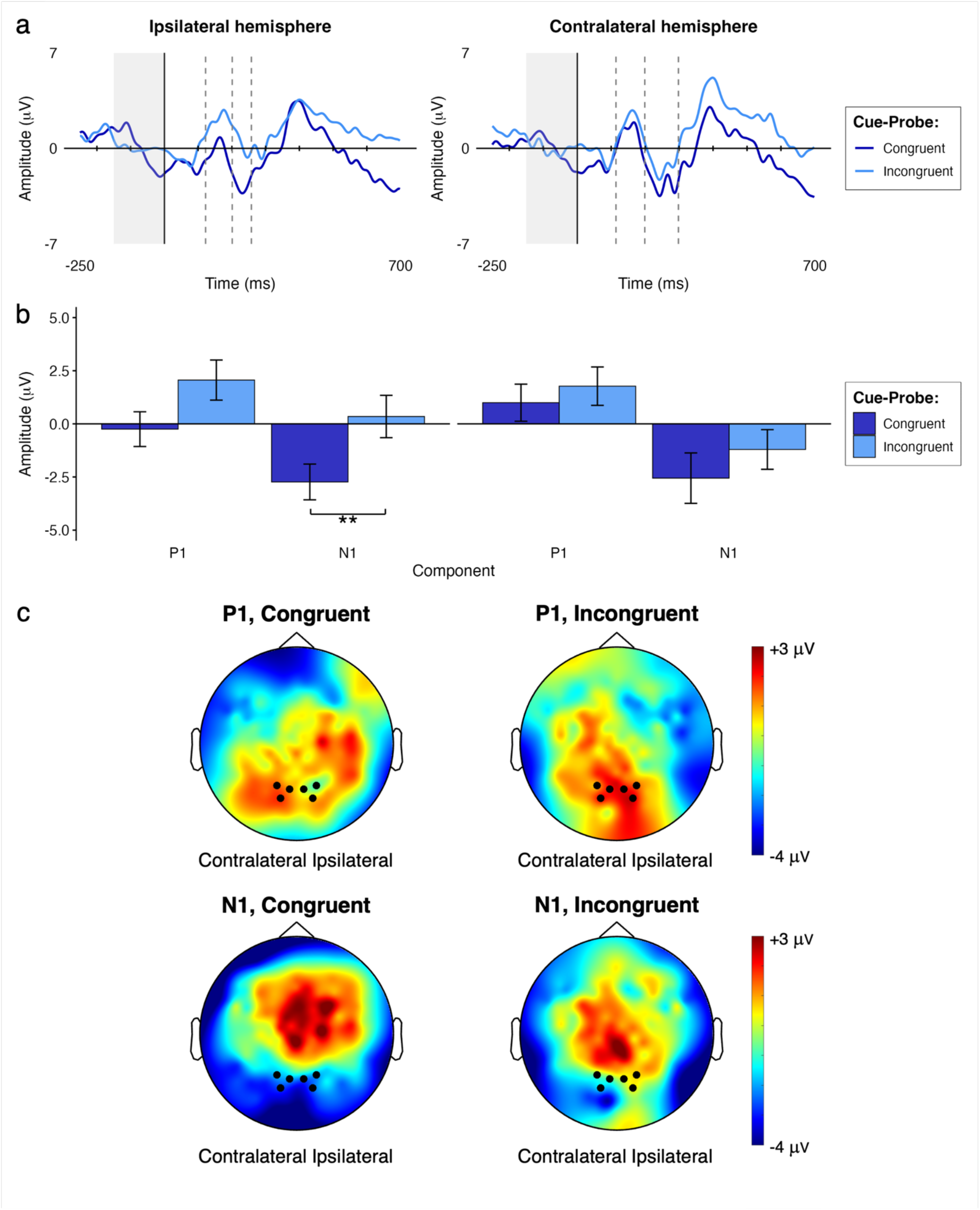
Modulation of 8-month-old infants’ VEPs by the congruency of the Cue-Probe pairing. **A.** Grand average VEPs in each hemisphere; the grey shading indicates the baseline period; dashed lines separate the time windows of components that were subjected to statistical comparison. **B.** Grand averaged mean individual amplitude of the VEPs in the two conditions for the two components of interest. **C.** Grand average topographical representations of the voltage distribution over the scalp in the two conditions (cue-probe Congruent and Incongruent) during the P1 and N1 components, with a Congruent – Incongruent difference map to the right; channels averaged for the analyses are highlighted (66, 70, 71 in the left hemisphere and 76, 83, 84 in the right hemisphere).

**Table 1.** Results of the (two-tailed) comparisons of the mean individual average of the 8-month-old infants’ VEP amplitudes in the Congruent vs Incongruent Cue-Probe conditions for the P1 and N1 components. Kolmogorov-Smirnov *D* statistics examine deviations from normality in the distributions of the differences between conditions.

| Component | *Kolmogorov-Smirnov* |  | *t-test* |  |  |  |
| --- | --- | --- | --- | --- | --- | --- |
|  | *D value* | *p value* | *dfs* | *t value* | *p value* | *d<sub>z</sub>* |
| Ipsilateral P1 | 0.161 | 0.653 | 18 | -1.740 | 0.099 | 0.399 |
| Ipsilateral N1 | 0.165 | 0.622 | 18 | -2.501 | 0.022 | 0.574 |
| Contralateral P1 | 0.211 | 0.320 | 18 | -0.745 | 0.466 | 0.171 |
| Contralateral N1 | 0.198 | 0.396 | 18 | -0.834 | 0.415 | 0.191 |

### 5-month-olds

As before we compared the mean individual amplitude of the VEPs in the Congruent vs Incongruent Cue-Probe conditions, in the P1 and N1 components, for both the Ipsilateral and the Contralateral hemispheres. The latencies of the components were: Ipsilateral hemisphere, P1: 110-212 ms, peaking at 190 ms; N1: 214-294 ms, peaking at 258 ms; Contralateral hemisphere, P1: 108-204 ms, peaking at 164 ms; N1: 206-330 ms, peaking at 252 ms (Fig. 2a).

**Figure 2.**
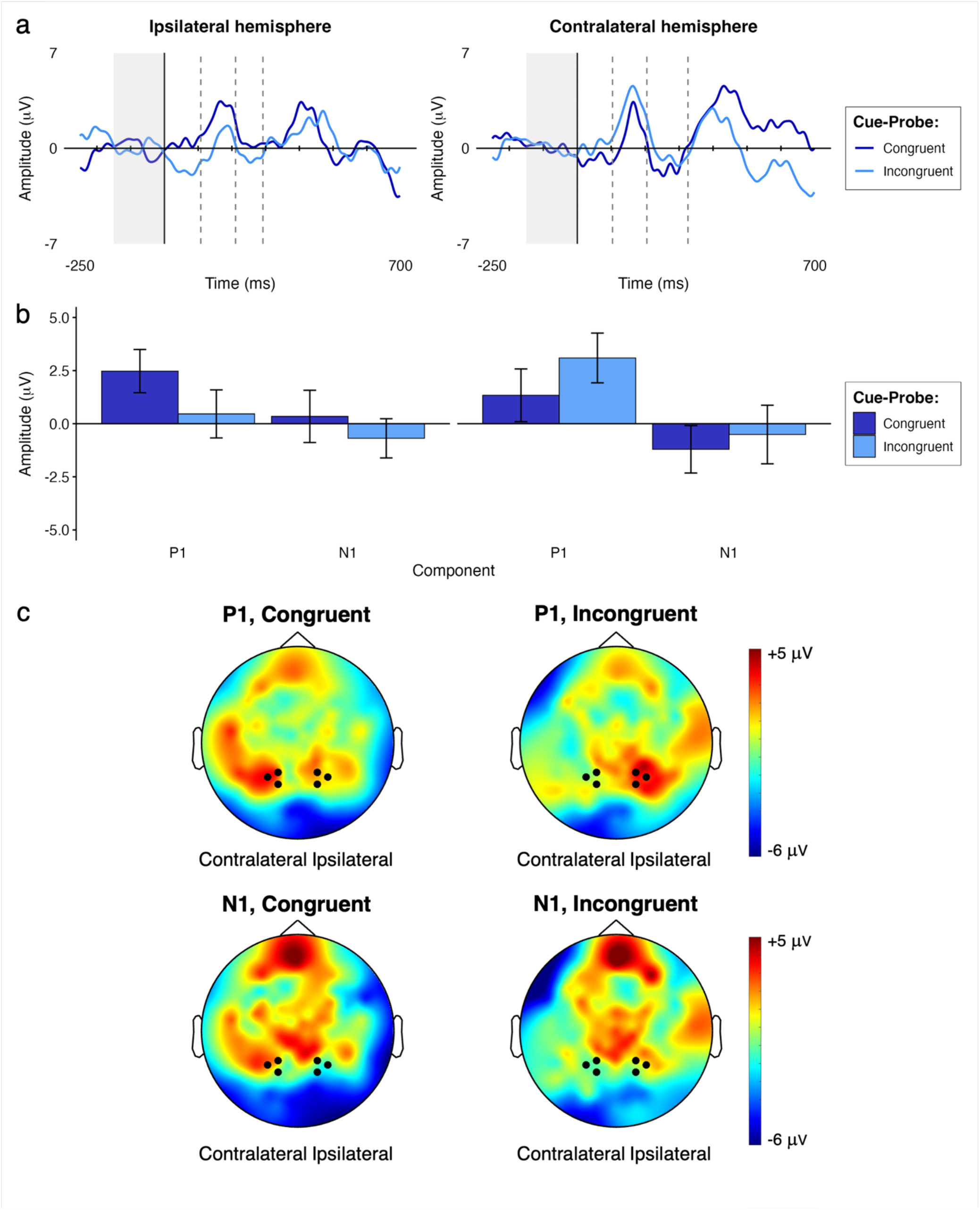
Modulation of 5-month-old infants’ VEPs by the congruency of the Cue-Probe pairing. **A.** Grand average VEPs in each hemisphere; the grey shading indicates the baseline period; dashed lines separate the time windows of components that were subjected to statistical comparison. **B.** Grand averaged mean individual amplitude of the VEPs in the two conditions for the two components of interest. **C.** Grand average topographical representations of the voltage distribution over the scalp in the two conditions (cue-probe Congruent and Incongruent) during the P1 and N1 components, with a Congruent – Incongruent difference map to the right; channels averaged for the analyses are highlighted (59, 60, 66 in the left hemisphere and 84, 85, 91 in the right hemisphere).

For each hemisphere and component, we ran a paired t-test to compare the mean individual amplitude of the waveform between the two Cue-Probe conditions (Fig. 2b). None of the comparisons reached statistical significance (Table 2).

**Table 2.** Results of the comparisons (two-tailed) of the mean individual average of the 5-month-old infants’ VEP amplitudes in the Congruent vs Incongruent Cue-Probe conditions for the P1 and N1 components. Kolmogorov-Smirnov *D* statistics examine deviations from normality in the distributions of the differences between conditions.

| Component | *Kolmogorov-Smirnov* |  | *t-test* |  |  |  |
| --- | --- | --- | --- | --- | --- | --- |
|  | *D value* | *p value* | *dfs* | *t value* | *p value* | *d<sub>z</sub>* |
| Ipsilateral P1 | 0.124 | 0.899 | 18 | 1.132 | 0.273 | 0.260 |
| Ipsilateral N1 | 0.168 | 0.601 | 18 | 0.599 | 0.557 | 0.137 |
| Contralateral P1 | 0.128 | 0.877 | 18 | -0.925 | 0.367 | 0.212 |
| Contralateral N1 | 0.094 | 0.990 | 18 | -0.330 | 0.745 | 0.076 |

## DISCUSSION

We presented 5- and 8-month-old participants with vibrotactile stimuli on one of their hands, followed by visual stimuli on the same (Congruent) or the opposite (Incongruent) hand, or no touch at all. The VEPs show that covert crossmodal tactile-visual spatial orienting is present at 8 months of age, while we found no evidence of crossmodal orienting at 5 months of age. More specifically, in the 8month-olds we observed an enhanced VEP in the first negative component after the presentation of the visual probe, but only when both the tactile and visual stimuli were presented on the same (vs different) hand, and only in the hemisphere ipsilateral to the hand to which the visuotactile stimulation was delivered. This mirrors findings observed in adult ERPs in a comparable tactile-visual crossmodal presentation (Kennett et al., 2001) where enhancements of the N1 following a congruent tactile cue were also seen ipsilaterally.

One interpretation of the modulation of 8-month-olds’ ipsilateral VEPs is that this demonstrates integration of the tactile and visual stimuli, rather than a crossmodal orienting effect. However, we find this interpretation unlikely given that the crossmodal interactions observed here in 8-month-olds are more indicative of crossmodal (tactile) cuing effects on visual processing that tend to be observed in early (typically within the first 300 ms in adults, Kennett et al., 2001; Ozawa & Yoshimura, 2024), occipital (O1 and O2) features of the VEP (P1 and N1) rather than of signatures of multisensory integration, which are not typically observed until the later stages of the visual processing hierarchy (Quinn et al., 2014). The observation of the ipsilateral N1 effect in common with Kennett et al. (2001) also points towards an interpretation in terms of crossmodal cuing.

The absence of a tactile-visual cuing effect in 5-month-old infants’ VEPs indicates that exogenous crossmodal spatial attention is undergoing development between 5 and 8 months of age. One possible interpretation is that 5 months of age is prior to the development of any capacity for attentional links between touch and vision. Perhaps more likely, 5-month-old infants’ crossmodal orienting may be at an earlier stage of development. Indeed, whilst none of the 5-month-olds’ comparisons were statistically reliable, the comparison of Congruent and Incongruent cue conditions in comparison in the ipsilateral P1 approached significance, highlighting perhaps an emerging substrate of crossmodal attention. One possibility is that tactile-visual cuing is present at this age but takes place more slowly. Further studies increasing the stimulus-onset-asynchronies of tactile cue and visual probe may reduce the speed-of-processing demands required of younger participants, enabling them to demonstrate crossmodal cuing. A further possibility is that the neural machinery supporting tactile-visual attention undergoes development between 5 and 8 months of age. Interestingly, the directions in which several components are modulated change between age groups. The amplitude of the ipsilateral P1 is larger for the Congruent condition in 5-month-olds but the converse is true for 8-month-olds; the amplitude of the ipsilateral N1 is more negative for the Incongruent condition in 5-month-olds with the converse true in 8-month-olds. Similar age-related differences are seen in the contralateral VEPs, suggesting the development of the brain functions supporting infants’ crossmodal cuing of visual attention.

The period of development between 5 and 8 months of age spans several salient changes in the ways in which the tactile and visual systems are coordinated. Overt crossmodal orienting of the visual system (the eyes) towards tactile stimuli on the hands emerges (Bremner et al., 2008) alongside visual spatial influences on tactile localisation (Begum Ali et al., 2015). Gori, Amadeo, and Bremner (Gori et al., in press), drawing on findings from blind infants and adults, have recently argued that these developments are driven by multisensory experience, and in particular the changing bodily dynamics affecting how visual, auditory and tactile systems interact in early life. A similar explanation can be applied to developments in tactile-visual crossmodal attention. With the developing coordination of hands and vision in the exploration of near space, infants gain increasingly organised and specific experiences of crossmodal contingencies between tactile and visual events, experiences that may support visual spatial attention (well established by 6 months of age) to become increasingly coordinated with tactile events and cues. The development of tactile-visual attention may also support the development of increasingly sophisticated sensorimotor action schemas. Further developmentally sensitive studies will be needed to fully disentangle the how crossmodal attention and sensorimotor functions interact in development.

The provision of multiple sensory modalities for perceiving the world and oneself is often presented as an adaptive boon for adults and infants alike, supporting advantages in precision, speed and accuracy (Stein, 2012, p. 2). And yet, from William James onwards, developmental scientists have pointed to the potential for confusion as infants attempt to make the right links between sensations arising in different senses (Bremner et al., 2012). This study sheds some critical light on this matter. The developmental changes in crossmodal attention between 5 and 8 months of age reported here show that the ability to shift visual attention towards the potential source of a tactile sensation is immature in early infancy, but bears a striking resemblance to adult crossmodal processing by 8 months of age. Furthermore, the current study introduces a task paradigm which can be used to investigate crossmodal attention across multiple time-frames and senses more widely, to inform our understanding of how infants develop their abilities to shift attention around their multisensory worlds.

## Supporting information

Supplementary results

## Author note

### Data availability

The present study has not been formally preregistered. The datasets generated and/or analysed during the current study are available in the University of Birmingham eData repository (Orioli & Bremner, 2025) and can be retrieved from https://doi.org/10.25500/edata.bham.00001374. The scripts and datasets used to perform the analyses reported in this manuscript are available online on the Open Science Framework website and can be retrieved from https://osf.io/k8qpe/.

## Acknowledgements

We are grateful to our participants and their parents for their invaluable contribution.

This research was funded by the European Research Council (FP7/2007-2013) (Grant agreement no. 241242 awarded to A.B.) and supported by the Leverhulme Trust (Early Career Fellowship ECF-2019-563 awarded to G.O).

## Author Contributions

A.B., R.T., J.B.A., and L.M. conceived the study and designed it. G.O., R.T., and J.B.A. collected the data and undertook data processing. A.B., G.O. and J.v.V. conducted the analyses. A.B. and G.O. wrote the report.

## Ethical approval

The parents were informed about the procedure and provided informed consent for their child’s participation. Ethical approval was gained from the Institutional Ethics Committee, Goldsmiths, University of London, and the study was conducted in accordance with the Declaration of Helsinki.

## REFERENCES

Azañón, E., Camacho, K., Morales, M., & Longo, M. R. (2018). The Sensitive Period for Tactile Remapping Does Not Include Early Infancy. Child Development, 89(4), 1394–1404. 10.1111/cdev.12813

Bahrick, L. E., & Lickliter, R. (2000). Intersensory redundancy guides attentional selectivity and perceptual learning in infancy. Developmental Psychology, 36(2), 190–201. 10.1037/0012-1649.36.2.190

Bahrick, L. E., Lickliter, R., & Flom, R. (2004). Intersensory Redundancy Guides the Development of Selective Attention, Perception, and Cognition in Infancy. Current Directions in Psychological Science, 13(3), 99–102. 10.1111/j.0963-7214.2004.00283.x

Bahrick, L. E., Lickliter, R., & Todd, J. T. T. (2020). The development of multisensory attention skills: Individual differences, developmental outcomes, and applications. In The Cambridge handbook of infant development: Brain, behavior, and cultural context (pp. 303–338). Cambridge University Press. 10.1017/9781108351959.011

Begum Ali, J., Spence, C., & Bremner, A. J. (2015). Human infants’ ability to perceive touch in external space develops postnatally. Current Biology, 25(20), R978–R979. 10.1016/j.cub.2015.08.055

Bremner, A. J., Lewkowicz, D. J., & Spence, C. (2012). Multisensory Development. SIPRI/Oxford University Press. 10.1093/acprof:oso/9780199586059.001.0001

Bremner, A. J., Lewkowicz, D. J., & Spence, C. (Eds.). (2012). Multisensory development (pp. x, 382). Oxford University Press. 10.1093/acprof:oso/9780199586059.001.0001

Bremner, A. J., Mareschal, D., Lloyd-Fox, S., & Spence, C. (2008). Spatial localization of touch in the first year of life: Early influence of a visual spatial code and the development of remapping across changes in limb position. Journal of Experimental Psychology: General, 137(1), 149–162. 10.1037/0096-3445.137.1.149

Butterworth, G., & Castillo, M. (1976). Coordination of auditory and visual space in newborn human infants. Perception, 5(2), 155–160. 10.1068/p050155

Chen, Y.-C., Lewis, T. L., Shore, D. I., Spence, C., & Maurer, D. (2018). Developmental changes in the perception of visuotactile simultaneity. Journal of Experimental Child Psychology, 173, 304–317. 10.1016/j.jecp.2018.04.014

Christodoulou, J., Leland, D. S., & Moore, D. S. (2018). Overt and covert attention in infants revealed using steady-state visually evoked potentials. Developmental Psychology, 54(5), 803–815. 10.1037/dev0000486

Clifton, R. K., Morrongiello, B. A., Kulig, J. W., & Dowd, J. M. (1981). Newborns’ orientation toward sound: Possible implications for cortical development. Child Development, 52(3), 833–838. 10.2307/1129084

Colombo, J. (2001). The development of visual attention in infancy. Annual Review of Psychology, 52, 337–367. 10.1146/annurev.psych.52.1.337

De Haan, M. (Ed.). (2013). Infant EEG and Event-Related Potentials. Psychology Press. 10.4324/9780203759660

Durand, K., Schaal, B., Goubet, N., Lewkowicz, D. J., & Baudouin, J.-Y. (2020). Does any mother’s body odor stimulate interest in mother’s face in 4-month-old infants? Infancy, 25(2), 151–164. 10.1111/infa.12322

Faul, F., Erdfelder, E., Lang, A.-G., & Buchner, A. (2007). G*Power 3: A flexible statistical power analysis program for the social, behavioral, and biomedical sciences. Behavior Research Methods, 39(2), 175–191. 10.3758/BF03193146

Gori, M., Amadeo, M. B., & Bremner, A. J. (In press). The development of multisensory space perception with and without early visual experience. Nature Reviews Psychology.

Hoehl, S., & Wahl, S. (2012). Recording Infant ERP Data for Cognitive Research. Developmental Neuropsychology, 37(3), 187–209. 10.1080/87565641.2011.627958

Hood, B. M. (1993). Inhibition of return produced by covert shifts of visual attention in 6-month-old infants. Infant Behavior and Development, 16(2), 245–254. 10.1016/0163-6383(93)80020-9

Hood, B. M., & Atkinson, J. (1993). Disengaging visual attention in the infant and adult. Infant Behavior & Development, 16(4), 405–422. 10.1016/0163-6383(93)80001-O

Hunt, A. R., & Kingstone, A. (2003a). Covert and overt voluntary attention: Linked or independent? Cognitive Brain Research, 18(1), 102–105. 10.1016/j.cogbrainres.2003.08.006

Hunt, A. R., & Kingstone, A. (2003b). Inhibition of return: Dissociating attentional and oculomotor components. Journal of Experimental Psychology: Human Perception and Performance, 29(5), 1068–1074. 10.1037/0096-1523.29.5.1068

James, W. (1890). The principles of psychology, Vol I. (pp. xii, 697). Henry Holt and Co. 10.1037/10538-000

Johnson, M. H. (1994). 11 Visual Attention and the Control of Eye Movements in Early Infancy. Attention and Performance XV: Conscious and Nonconscious Information Processing, 15, 291.

Johnson, M. H., Posner, M. I., & Rothbart, M. K. (1994). Facilitation of Saccades Toward a Covertly Attended Location in Early Infancy. Psychological Science, 5(2), 90–93. 10.1111/j.1467-9280.1994.tb00636.x

Johnson, M. H., & Tucker, L. A. (1996). The development and temporal dynamics of spatial orienting in infants. Journal of Experimental Child Psychology, 63(1), 171–188. 10.1006/jecp.1996.0046

Kennett, S., Eimer, M., Spence, C., & Driver, J. (2001). Tactile-Visual Links in Exogenous Spatial Attention under Different Postures: Convergent Evidence from Psychophysics and ERPs. Journal of Cognitive Neuroscience, 13(4), 462–478. 10.1162/08989290152001899

Leleu, A., Rekow, D., Poncet, F., Schaal, B., Durand, K., Rossion, B., & Baudouin, J.-Y. (2020). Maternal odor shapes rapid face categorization in the infant brain. Developmental Science, 23(2), e12877. 10.1111/desc.12877

Luck, S. J., & Gaspelin, N. (2017). How to get statistically significant effects in any ERP experiment (and why you shouldn’t). Psychophysiology, 54(1), 146–157. 10.1111/psyp.12639

Moreau, T., Helfgott, E., Weinstein, P., & Milner, P. (1978). Lateral differences in habituation of ipsilateral head-turning to repeated tactile stimulation in the human newborn. Perceptual and Motor Skills, 46(2), 427–436. 10.2466/pms.1978.46.2.427

Muir, Clifton, R. K., & Clarkson, M. G. (1989). The development of a human auditory localization response: A U-shaped function. Canadian Journal of Psychology / Revue Canadienne de Psychologie, 43(2), 199–216. 10.1037/h0084220

Muir, & Field, J. (1979). Newborn infants orient to sounds. Child Development, 50(2), 431–436. 10.2307/1129419

Oostenveld, R., Fries, P., Maris, E., & Schoffelen, J.-M. (2011). FieldTrip: Open Source Software for Advanced Analysis of MEG, EEG, and Invasive Electrophysiological Data. Computational Intelligence and Neuroscience, 2011, 1–9. 10.1155/2011/156869

Orioli, G., & Bremner, A. J. (2025). Cortical signatures of crossmodal attention in the first year of life. [Data set]. uBIRA. 10.25500/edata.bham.00001374

Orioli, G., Bremner, A. J., & Farroni, T. (2018). Multisensory perception of looming and receding objects in human newborns. Current Biology, 28(22), R1294–R1295. 10.1016/j.cub.2018.10.004

Orioli, G., Parisi, I., Van Velzen, J. L., & Bremner, A. J. (2023). Visual objects approaching the body modulate subsequent somatosensory processing at 4 months of age. Scientific Reports, 13(1), 19300. 10.1038/s41598-023-45897-4

Ozawa, Y., & Yoshimura, N. (2024). Temporal Electroencephalography Traits Dissociating Tactile Information and Cross-Modal Congruence Effects. Sensors, 24(1), 45. 10.3390/s24010045

Popov, T., Oostenveld, R., & Schoffelen, J. M. (2018). FieldTrip Made Easy: An Analysis Protocol for Group Analysis of the Auditory Steady State Brain Response in Time, Frequency, and Space. Frontiers in Neuroscience, 12, 711. 10.3389/fnins.2018.00711

Posner, M. I. (1980). Orienting of attention. The Quarterly Journal of Experimental Psychology, 32(1), 3–25. 10.1080/00335558008248231

Posner, M. I., Rafal, R. D., Choate, L. S., & Vaughan, J. (1985). Inhibition of return: Neural basis and function. Cognitive Neuropsychology, 2(3), 211–228. 10.1080/02643298508252866

Quinn, B. T., Carlson, C., Doyle, W., Cash, S. S., Devinsky, O., Spence, C., Halgren, E., & Thesen, T. (2014). Intracranial Cortical Responses during Visual–Tactile Integration in Humans. Journal of Neuroscience, 34(1), 171–181. 10.1523/JNEUROSCI.0532-13.2014

Richards, J. E. (1987). Infant visual sustained attention and respiratory sinus arrhythmia. Child Development, 58(2), 488–496.

Richards, J. E. (1997). Effects of attention on infants’ preference for briefly exposed visual stimuli in the paired-comparison recognition-memory paradigm. Developmental Psychology, 33(1), 22–31. 10.1037//0012-1649.33.1.22

Richards, J. E. (2000). Localizing the development of covert attention in infants with scalp eventrelated potentials. Developmental Psychology, 36(1), 91–108. 10.1037/0012-1649.36.1.91

Rigato, S., Banissy, M. J., Romanska, A., Thomas, R., Van Velzen, J., & Bremner, A. J. (2019). Cortical signatures of vicarious tactile experience in four-month-old infants. Developmental Cognitive Neuroscience, 35, 75–80. 10.1016/j.dcn.2017.09.003

Rochat, P. (2021). Clinical pointers from developing self-awareness. Developmental Medicine & Child Neurology, 63(4), 382–386. 10.1111/dmcn.14767

Schaal, B., & Durand, K. (2012). The role of olfaction in human multisensory development. In Multisensory development (pp. 29–62). Oxford University Press. 10.1093/acprof:oso/9780199586059.003.0002

Slater, A., Quinn, P. C., Brown, E., & Hayes, R. (1999). Intermodal perception at birth: Intersensory redundancy guides newborn infants’ learning of arbitrary auditory−visual pairings. Developmental Science, 2(3), 333–338. 10.1111/1467-7687.00079

Spence, C., & Driver, J. (1994). Covert spatial orienting in audition: Exogenous and endogenous mechanisms. Journal of Experimental Psychology: Human Perception and Performance, 20(3), 555–574. 10.1037/0096-1523.20.3.555

Spence, C., & Driver, J. (1996). Audiovisual links in endogenous covert spatial attention. Journal of Experimental Psychology: Human Perception and Performance, 22(4), 1005–1030. 10.1037/0096-1523.22.4.1005

Spence, C., & Driver, J. (1997). On measuring selective attention to an expected sensory modality. Perception & Psychophysics, 59(3), 389–403. 10.3758/BF03211906

Spence, C., & Driver, J. (2004). Crossmodal space and crossmodal attention. Oxford University Press.

Spence, C., Nicholls, M. E. R., Gillespie, N., & Driver, J. (1998). Cross-modal links in exogenous covert spatial orienting between touch, audition, and vision. Perception & Psychophysics, 60(4), 544–557. 10.3758/BF03206045

Stein, B. E. (Ed.). (2012). The New Handbook of Multisensory Processing. MIT Press.

Webb, S. J., Long, J. D., & Nelson, C. A. (2005). A longitudinal investigation of visual event-related potentials in the first year of life. Developmental Science, 8(6), 605–616. 10.1111/j.1467-7687.2005.00452.x

Wertheimer, M. (1961). Psychomotor coordination of auditory and visual space at birth. Science (New York, N.Y.), 134(3491), 1692. 10.1126/science.134.3491.1692

Xie, W., & Richards, J. E. (2017). The Relation between Infant Covert Orienting, Sustained Attention and Brain Activity. Brain Topography, 30(2), 198–219. 10.1007/s10548-016-0505-3

