## Supplementary results for "Crossmodal attention develops in the first year of life: Cortical signatures of tactile to visual exogenous spatial cuing at 8 but not 5 months of age"

[†] current affiliations: A.B. and G.O., Centre for Developmental Science, School of Psychology, University of Birmingham (Birmingham, United Kingdom); R.T., University College London (London, UK); J.B.A., Centre for Brain and Cognitive Development, Faculty of Science, Department of Psychological Sciences, Birkbeck, University of London (London, UK); L.M, [5]Department of Forensic and Neurodevelopmental Sciences, Institute of Psychiatry, Psychology, and Neuroscience, King's College London (London, UK).

### RESULTS

#### 5-month-olds

*Results of the t-tests after having removed 3 participants for which the behavioural coding of the videos is not available*

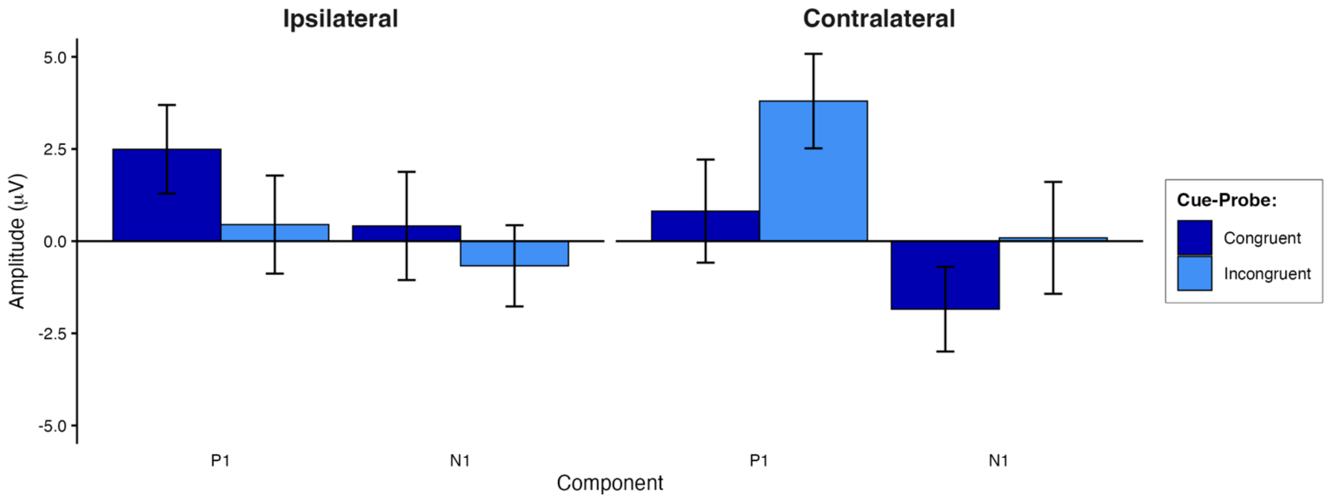


**Supplementary Figure SI1.** Voltage differences in the grand averaged mean individual amplitude of 5-month-olds infants' VEPs in the two conditions for the two components of interest. Only infants for whom the video recordings were available are included (n = 16).

**Supplementary Table SI1.** The table summarises the results of the comparisons (two-tailed) of the mean individual average of the 5-month-old infants' VEP amplitudes in the Congruent vs Incongruent Cue-Probe conditions for the P1 and N1 components. Only infants for whom the video recordings were available are included (n = 16). Kolmogorov-Smirnov D statistics identifying significant deviations from normality in the distributions of the differences between conditions are also included.

| **Component** | **Kolmogorov-Smirnov** | | **t-test** | | | |
|---|---|---|---|---|---|---|
| | **D value** | **p value** | **dfs** | **t value** | **p value** | **$d_z$** |
| Ipsilateral P1 | 0.138 | 0.883 | 15 | 0.965 | 0.35 | 0.241 |
| Ipsilateral N1 | 0.216 | 0.390 | 15 | 0.526 | 0.607 | 0.132 |
| Contralateral P1 | 0.143 | 0.854 | 15 | -1.459 | 0.165 | 0.365 |
| Contralateral N1 | 0.097 | 0.994 | 15 | -0.869 | 0.399 | 0.217 |

### METHOD

#### EEG recording and analyses

**Supplementary Table SI2.** Mean number of artefact-free trials for the 5-month-old age group, condition and trial type (C stands for Cue, P for Probe, L for Left and R for Right).

| measure | CL-PL | CL-PR | CL-No P | CR-PL | CR-PR | CR-No P | Congruent | Incongruent | No Probe |
|---|---|---|---|---|---|---|---|---|---|
| **mean** | 14.16 | 13.68 | 13.79 | 14.16 | 14.26 | 14.79 | 28.42 | 27.84 | 28.58 |
| **S.D.** | 4.57 | 4.47 | 4.76 | 4.49 | 4.76 | 5.91 | 8.98 | 8.34 | 10.17 |
| **min** | 7.00 | 8.00 | 7.00 | 9.00 | 7.00 | 8.00 | 15.00 | 18.00 | 15.00 |
| **max** | 24.00 | 24.00 | 26.00 | 26.00 | 26.00 | 32.00 | 50.00 | 50.00 | 54.00 |

**Supplementary Table SI3.** Mean number of artefact-free trials for the 7-month-old age group, condition and trial type (C stands for Cue, P for Probe, L for Left and R for Right).

| measure | CL-PL | CL-PR | CL-No P | CR-PL | CR-PR | CR-No P | Congruent | Incongruent | No Probe |
|---|---|---|---|---|---|---|---|---|---|
| **mean** | 14.63 | 13.74 | 13.21 | 14.47 | 13.63 | 13.42 | 28.26 | 28.21 | 26.63 |
| **S.D.** | 5.56 | 6.39 | 6.82 | 5.27 | 4.95 | 6.65 | 10.02 | 11.29 | 13.06 |
| **min** | 6.00 | 5.00 | 6.00 | 5.00 | 7.00 | 4.00 | 13.00 | 12.00 | 11.00 |
| **max** | 27.00 | 25.00 | 30.00 | 25.00 | 25.00 | 26.00 | 52.00 | 49.00 | 56.00 |
